# Impact of protein degradation and cell growth on mammalian proteomes

**DOI:** 10.1101/2025.02.10.637566

**Authors:** Andrew Leduc, Shanshan Zheng, Purvi Saxena, Nikolai Slavov

**Affiliations:** Departments of Bioengineering, Biology, Chemistry and Chemical Biology, Single Cell Proteomics Center, and Barnett Institute, Northeastern University, Boston, MA 02115, USA; Parallel Squared Technology Institute, Watertown, MA 02472, USA

## Abstract

Cellular protein abundances are controlled by rates of synthesis and clearance. Since the latter includes protein degradation and dilution due to cell growth, the growth rate may influence the mechanisms controlling variation in protein abundances. To quantify this influence, we analyzed the growth-dependent effects of protein degradation within a cell type (between activated and resting human B-cells), across human cell types and mouse tissues. This analysis benefited from deep and accurate quantification of over 12,000 proteins across all four primary tissues using plexDIA. The results indicate that growth-dependent dilution accounts for 40 % of protein abundance changes across conditions. Furthermore, the variation in protein clearance rates is sufficient to explain up to 50 % of the protein abundance differences within slowly growing cells as contrasted with 7 % in growing cells. Remarkably, clearance rates also regulate the abundance differences of proteoforms encoded by the same gene, such those arising from alternative splicing or alternate RNA decoding. Overall, our results demonstrate substantially larger than previously appreciated contributions of protein degradation to protein variation at slow growth, both across proteoforms and tissue types.

## Introduction

Protein abundances vary across the proteome and across cell types and states. This variation is controlled by protein synthesis and clearance. Protein synthesis rates reflect both mRNA abundance and translation efficiency, while clearance rates include protein degradation, secretion, and dilution from cell growth. Synthesis rates have been shown to be the dominant factor in a few model systems representing growing cells^1–3^. These studies estimate that protein clearance explains just 5-8% of the variation in absolute abundance across the proteome. Yet, the relative importance of protein synthesis and clearance may depend on the rate of cell growth as hypothesized previously^3–5^.

At low growth rates, proteins accumulate according to their rate of degradation or secretion, producing an inverse relationship between clearance rate and protein abundance. As the growth rate increases, this accumulation is attenuated because proteins become diluted faster than they are degraded or secreted. As proteins accumulate depending on their rate of degradation or secretion, this attenuation will alter protein abundances unless synthesis or degradation rates change to compensate^4,5^. Although significant compensation has been demonstrated, it is not yet clear to what degree changes in protein synthesis and degradation fully counterbalance the shift from degradation-driven clearance to dilution-driven clearance.

Furthermore, the significance of this compensation, or lack there of, depends on the degree of variability in degradation and secretion rates. Simply put, if all proteins have relatively similar degradation and secretion rates, there would be minimal potential for preferential accumulation. To measure clearance as well as synthesis rates across the proteome, liquid chromatography tandem mass spectrometry (LC-MS/MS) combined with metabolic pulse labeling has become the tool of choice, both in vitro and in vivo^6^. Such measurements are greatly facilitated by recent advances, such as plexDIA^7,8^. Metabolic pulse studies have reported that protein specific degradation rates can span three orders of magnitude^1,3,9–12^. This suggests that changes in cell growth require protein synthesis to compensate substantially if protein abundances are to be maintained.

In addition to variation between different proteins, studies have reported significant differences in degradation rates for the same protein in different cell types^10^ and tissues^12^. Analysis in Hela cells has even reported that individual protein isoforms may have distinctive degradation rates^13^. However, the extent to which differences in protein degradation contribute to tissue and cell type specific protein and proteoform abundance is an open question demanding robust measurements and analysis incorporating reliability estimates. Further, the magnitude and scope of this regulatory impact remains unclear in primary cells and tissues.

In this study, we set out to explore these questions using measurements of protein abundance, synthesis and clearance, including different data acquisition approaches for reliability estimates. Using *in vivo* metabolic pulse labeling with lysine having heavy isotopes, different proteases and MS instruments, we acquired plexDIA datasets with high quantitative accuracy, consistency and proteome coverage. This includes protein abundances for over 12,000 proteins and degradation rates for over 10,000 proteins from over 113,000 lysine containing peptides, thus supporting proteoform quantification. These data strongly support our finding that the average rate of cell growth within tissue strongly predicts the amount of variation in protein abundances explained by clearance. Remarkably, we find that in slowly growing tissues such as the brain, protein degradation explains approximately half the variation in protein abundance. Further, we establish that differences in growth and degradation rates across cell states, cell types, and tissues substantially shape relative protein abundances. Having established that degradation strongly impacts the abundance of proteins from different genes, we explored whether degradation also impacts the abundance of proteoforms templated by the same gene. We quantified this impact for proteoforms originating from alternative splicing or from alternate RNA decoding^14^. The results demonstrate strong impact, extending the role of degradation to setting the abundance of proteoforms originating from the same gene.

## Results

### The impact of clearance on protein abundances varies across cell types

We first set out to explore factors that control the extent of absolute protein abundance regulation by protein clearance. To do so, we quantified protein clearance in a variety of cell types, including several that were not actively proliferating. To estimate the influence of protein clearance, we must first define its relationship to protein abundance. In non-growing cells, the steady-state abundance for the *i*th protein (*p_i_*) equals the ratio of the rates of protein synthesis *k_s,i_* and protein degradation or secretion *k_d,i_*, Fig. 1a. In growing cells, an additional term is needed to account for dilution of proteins due to the increase in volume that occurs as cells grow, *k_g_*^5,15^, Fig. 1a. Thus, protein abundance is inversely proportional to the overall amount of protein clearance *k_r,i_*, the combined influence of degradation, secretion, and dilution, Fig. 1a. As synthesis and clearance rates are well approximated by the log-normal distribution^16^, Fig. EV1a, we define the protein abundance variation explained by clearance, *R*^2^, as the sum of squared residuals explained solely by *K_r_* in the following equation: log(*P*) = log(*K_s_*) *−* log(*K_r_*). This approach is more robust than the square of the Pearson correlation, which is agnostic of the regression coefficient and thus can overestimate when the rates of synthesis and degradation are correlated Fig. EV1b. It should be noted that log transforming the data places more uniform emphasis on the contribution of regulation to each protein, versus prioritizing total protein mass.

**Figure 1.**
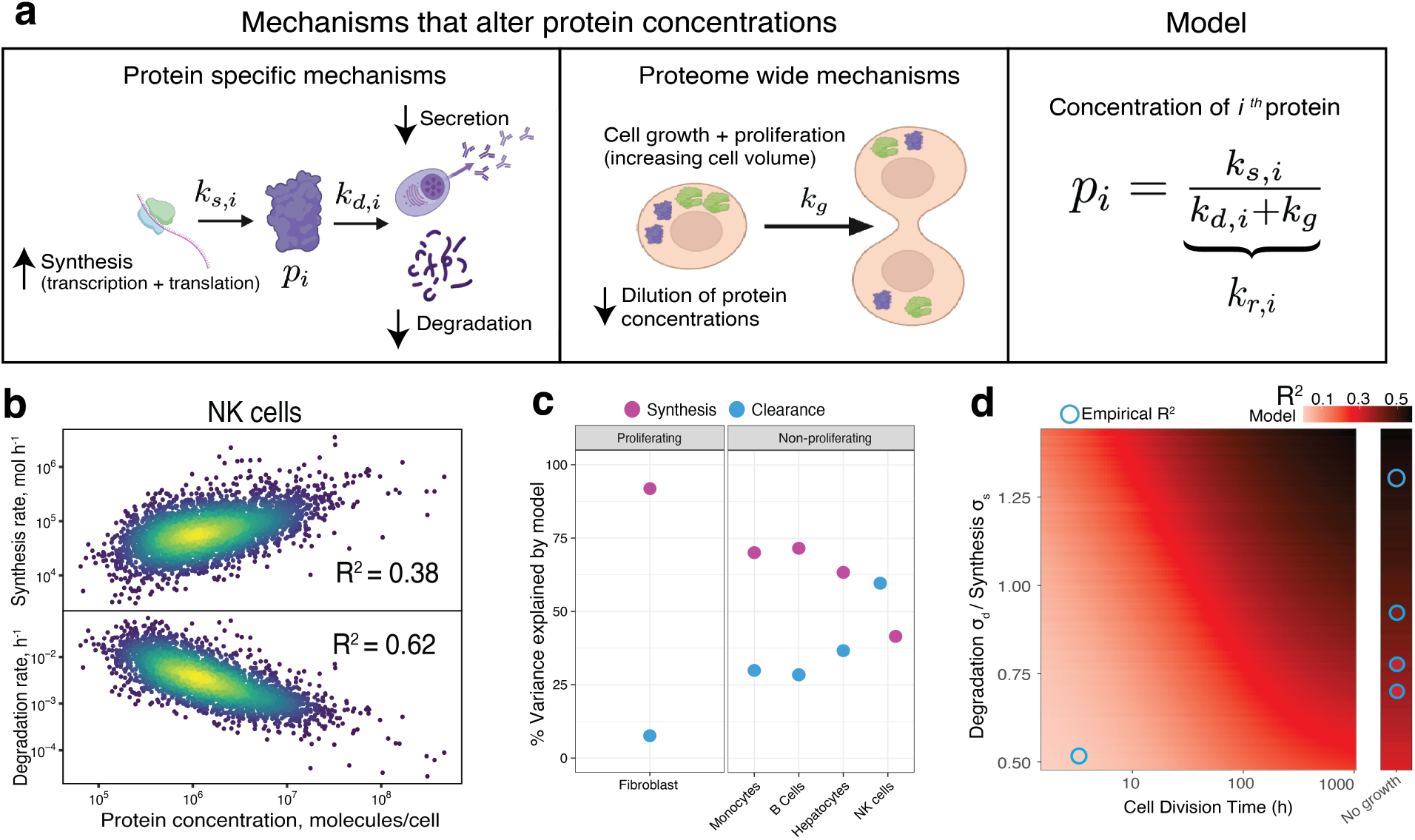
Modeling the impact of degradation on protein abundance in proliferating and non-proliferating human cells. **a**, Protein abundance (*p_i_*) is determined by protein specific factors such as synthesis (*k_s,i_*) and degradation / secretion rates (*k_d,i_*) as well as global factors such as the rate of dilution caused by cell growth (*k_g_*). **b**, Data from non-proliferative natural killer collected by Mathieson *et al.*^10^ show strong dependence of protein abundance on rates of degradation, *R*^2^ = 0.62, suggesting that protein degradation account for 62 % of the variation of protein abundances. Protein synthesis rates due to transcription and translation account for 38 % of the variation in protein abundances in natural killer cells. **c**, The % variance explained by degradation and synthesis varies for proliferating cells, fibroblasts collected by Schwanhauser *et al.*^1^, and additional non-growing cell types collected by Mathieson *et al.*^10^. **d**, Model predictions for the variance of protein abundances explained by degradation as a function of cell division time and the ratio of standard deviations for the log normally distributed rates of protein degradation (*k_s_*) and synthesis (*k_d_*). Predictions align with empirical estimates derived via regression.

Applying this approach to metabolic pulse labeling LC-MS/MS data, we analyzed proliferative cells from Schwanhausser *et al.*^1^ and non-proliferative cell types from Mathieson *et al.*^10^ Remarkably, *R*^2^ estimates for the fraction of absolute abundance variation explained by clearance span a wide range, varying from 7 % in proliferative fibroblasts to 62 % in non-proliferative natural killer cells, Fig. 1b,c. Interpreting these estimates requires accounting for the influence of measurement noise. Systematic biases, such as ionization efficiency and digestion efficiency, cannot be accounted for with replicate measurements. These biases predominantly affect the estimation of protein abundance and synthesis rates. When estimating clearance rates, these biases similarly affect light and heavy peptides and thus cancel out when computing ratios. Because systematic biases are not shared between the estimates of protein clearance and abundance, they are unlikely to inflate our *R*^2^ estimates.

The contrast between clearance explaining just 7 % of absolute protein abundance in dividing cells to between 28% and 62% in non-dividing cells suggests growth rate is a significant factor. However, the variation across non-dividing cells suggests contribution of factors beyond cell growth as well. One possible factor is cell type specific rates of protein degradation and secretion^10^. Another possible factor is that although these cell types are not actively proliferating, culturing primary cells *in vitro* has been shown to cause changes to cell size^17^. These size changes may inflate or decrease the influence of protein clearance. These factors, as well as differences in measurement quality and depth of coverage, may also explain differences with reports of non-proliferative cells where only 8-11 % variance has been explained by protein clearance^3^.

These observations suggest a simple model that we sought to formalize so that we can simulate testable predictions for the effect of growth rate. Specifically, we modeled the synthesis and clearance rates from a bi-variate log-normal distribution. The extent of co-variation between synthesis and degradation was modest in the samples we analyzed, Fig. EV1c. We speculate that the negative correlation present only for proliferative fibroblasts may reflect the compensation in protein synthesis rates for long lived proteins, as increased synthesis is needed to make up for the decreased accumulation of proteins with slower degradation rates. However, because co-variation was modest, we chose to set the co-variation to 0 falling between estimates for growth and non-growth.

Using this model, protein abundances were then computed from the ratio of simulated synthesis and clearance rates, where clearance rates equal the degradation rate plus the growth rate dilution constant, Fig. 1a. We simulated data through a physiologically relevant range of growth rates and variances for the distribution of synthesis and degradation rates, Fig. EV1d. The resulting *R*^2^ estimates closely match empirical estimates across a wide range of parameters (Fig. EV1d), thus suggesting explanations to both questions raised from the results in Fig. 1c. First, we observed that degradation rates have cell-type specific degrees of variability. As a result, the influence of degradation differed across non-growing cell types, Fig. 1d and Fig. EV1d. Secondly, the results confirmed that the level of growth in the proliferative fibroblasts could indeed quantitatively explain the differing amounts of variance explained by degradation in growth and non-growth conditions Fig. 1d. The correspondence between model predication and empirical estimates provides some support for the model, but the data points are too few to establish confidence. Therefore, we set out to acquire more data and perform additional analysis testing the core principles of our model.

Whereas protein clearance rates have been measured in relatively few cell types, growth rates^18^, protein and transcript abundances have been measured across hundreds of cancer cell lines^19,20^. This offered an opportunity to test a key prediction of our model: that the control of protein abundance by transcription, an essential step of synthesis, should increase with growth rate as the contribution of clearance declines. We therefore assembled growth rates for the 375 cell lines profiled at both the proteomic and transcriptomic level. Because these growth rates were not measured in the same cultures used for the omic profiling, we first assessed how reproducible growth-rate measurements are across cultures. For the 982 lines with two or more independent reports, doubling times were moderately reproducible, Fig. EV2a, (Pearson *r* = 0.59). We then asked how the within-line correlation between absolute transcript and protein abundance scales with growth rate, and found a weak but significant positive relationship, (*r* = 0.13, *p* = 0.022) Fig. EV2b, consistent with transcriptional control increasing modestly with growth rate. To reduce the influence of noise in the literature growth rates, we next estimated proliferation directly from each transcriptome, using the mean expression of canonical cell-cycle markers as a proliferation signature derived from the same sample. This signature was more strongly associated with the transcript-protein correlation (*r* = 0.37, *p* = 3 *×* 10*^−^*^13^), Fig. EV2c. Together, these results provide additional support for an increasing contribution of protein synthesis to protein abundance as cell growth rate rises.

### Incomplete compensation for protein dilution upon B-cell activation

To test the dependence between growth rate and protein clearance in modeled in Fig. 1d, we next sought to contrast changes in growth more directly. To this end, we first focused on growth rate changes within a cell type, namely primary B-cells in the resting state (non proliferative) and activated state (highly proliferative). We utilized abundance measurements made by Rieckmann *et al.*^21^ and degradation rate measurements made in resting primary B-cells by Mathieson *et al.*^10^. This system was ideal for several reasons. First, without stimulation, B cells can persist for months in vivo without dividing^22^. However, upon stimulation, B-cells can divide as rapidly as every 16 hours, providing a large growth rate differential^23^, Fig. 2a. Additionally, degradation rates are likely to be more similar within as opposed to across cell-type. Thus, measuring changes within a cell type provides a relatively controlled comparison. Additionally, replicate measurements of protein abundance and protein clearance allowed us to estimate and account for random noise in the estimation of protein fold changes and protein clearance, see methods.

**Figure 2.**
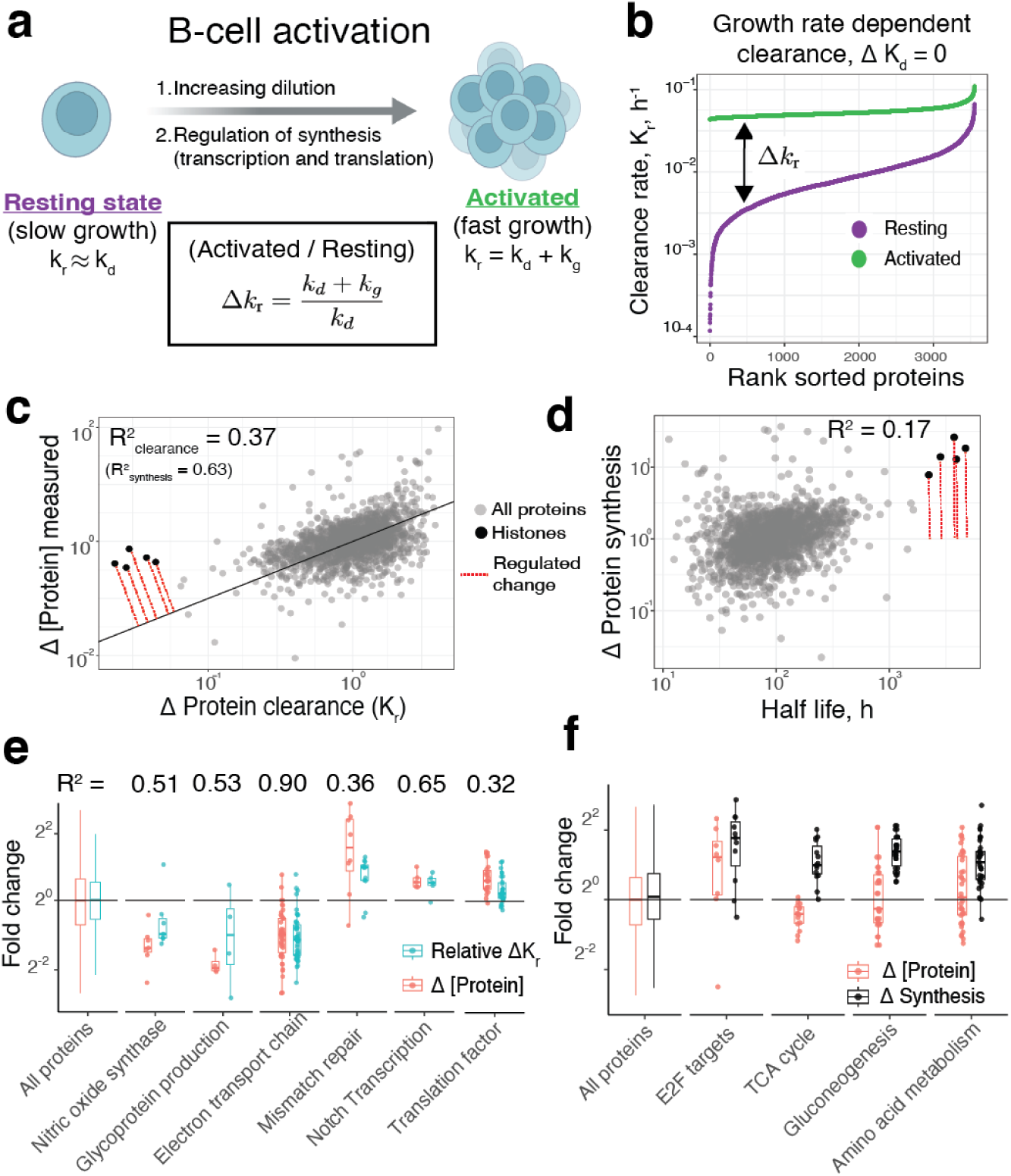
Proteome differences between resting and activated B-cells originate from altered protein dilution. **a**, When B cells are exposed to antigen, they transition from a resting state where there is minimal cell growth, to an activated state where cells divide every 16 hours^5^. **b**, Thus, clearance rates increase for all proteins with *k_g_*, resulting in larger relative changes for more stable proteins. **c**, The relative changes in clearance rates between the activated and resting state explains 37 % of the variance in protein abundance changes between these two states. **d**, However, not all proteins change abundance proportional to the shift in clearance. Many proteins such as histones are actively regulated with increased protein synthesis tied to the rate of cell division.**e**, Functional protein groups driven by dilution include the electron transport chain where 90 % of protein fold changes are caused by increased protein clearance due to growth. **f**, Proteins involved in processes such as gluconeogenesis get regulated to increase their abundance to counteract the increased rate of clearance.

While others have emphasized the regulated changes in gene expression, transcription and translation, that compensate for growth related changes in protein abundance^4,5^, our model quantifies protein abundance changes resulting solely from increased dilution caused by cell growth. In this model, as growth rate increases, the production of all proteins must increase. We define this increase uniformly for each protein as a constant *δ*. Deviations in synthesis from the expected scalar for a given protein *i* are defined with an additional term *µ_i_*. Thus the change in synthesis upon an increase in growth rate can be defined as Δ*k_s,i_* = *δµ_i_*. If *δ* explains all increases in protein synthesis, *µ_i_*= 1 for all proteins, changes in clearance caused by dilution will alter protein abundances. To demonstrate why, in non-growing B cells, protein clearance equals protein degradation rate and spans a dynamic range from 10*^−^*^4^ to 10*^−^*^1^, Fig. 2b. As uniform protein dilution outpaces degradation in activated B cells, clearance rates span only 10-fold dynamic range, Fig. 2b. Thus, with only a uniform scalar change in synthesis rate, protein abundances will decrease upon activation proportionally to the change in clearance, Fig. 2b. Again, this scenario represents an absence of compensation.

When comparing abundance changes and expected clearance changes, we find that change in clearance caused by protein dilution alone explains 37 % of the variation in protein abundances between the activated and resting state, Fig. 2c. This demonstrates that compensation incompletely accounts for a significant proportion of changes in protein clearance. However, proteins such as histones notably defy this trend and maintain their abundance. In these cases, synthesis rates are regulated to compensate and prevent significant abundance decrease. We interpret these deviations from the expected growth induced dilution as regulated changes in synthesis, *µ_i_*. Regulated changes correlate more weakly with protein half life explaining only 17 % of the variation in regulated changes. This suggests that the majority of changes in synthesis do not specifically work to counteract dilution, Fig. 2d.

Note that our minimal model for clearance changes upon B-cell activation focuses only on changes caused by growth increase. Because dilution dominates degradation at high growth, changes in degradation rates associated with activation will have a diminished effect on changes in clearance, except potentially for the least stable proteins. Further, any substantial changes in degradation would reduce the explanatory power of our model, which assumes that degradation rates remain constant between conditions. Thus, our model will report conservative estimates for the impact of growth induced clearance changes. Further, note that measurements come from different labs and variation between the B cells, such as donor and culture protocol, will result in misaligned degradation rates and protein fold change estimates. We believe, this will lead to an underestimation of percent variance explained by our model as these differences would attenuate the trend. However, unknown systematic effects that could introduce artificially inflated results cannot be excluded.

To this end, we wanted to confirm our results in a system where clearance rates were obtained in both activated and resting cells from the same lab. We turned to data from Wolf et al. where resting and activated T-Cells from the same donor were treated with pulse SILAC^24^. First, we tested whether the same general relationship was present and found trends similar to the B cells: protein fold changes between activated and resting T Cells demonstrated half-life dependent dilution (Fig. EV3a) and proteins with longer half lives have lower abundance in activated cells (Fig. EV3b), *r* = *−*0.21. The slightly weaker trend observed in activated T Cells may be due to the a slower growth rate for this specific experiment. When comparing the clearance half lives between resting and activated T Cells, the difference in slope suggests that these cells had a 31 hour doubling time, roughly twice as slow as the 16 hours reported for the B cells from Rieckmann et al.^21^. Importantly, we did not observe a systematic shift in clearance rates from the activated T-cells that would counter act the dilution. Using the difference in activated versus resting clearance rates modestly increased the explanatory power of the activated vs resting fold changes, *r* = *−*0.35.

Because dilution induced abundance changes are largely not compensated for, this raises the question of whether this dilution contributes to defining the phenotype of the activated state. Towards this direction, proteins from a number of concerted pathways change their abundance substantially between the activated and resting state. Many of these pathways can attribute over half the variance or more to the changes in protein clearances caused by the change in dilution, Fig. 2e. For example, proteins participating in the electron transport chain decrease their abundance by two fold and are predominantly driven by dilution. This may reflect the decrease in metabolism from oxidative phosphorylation to glycolysis during activation^25^. Other proteins weakly affected by dilution such as translation factors and proteins participating in notch transcription increase in abundance, Fig. 2e. This may reflects the increased demand for protein synthesis. Conversely, examining the extent of active regulation reveals several pathways for which changes in synthesis rate exceed changes in protein abundance. These pathways include the E2F transcription pathway as well as proteins participating in gluconeogenesis, Fig. 2f. Regulation of metabolic enzymes participating in glucose metabolism may be required to meet the energy demands of increased proliferation. However, because this analysis rests on the intersection of two externally generated datasets, only *∼*1,200 proteins were quantified in common, which limited resolution of smaller or activation-specific functional categories.

### Growth and degradation rate differences shapes tissue-specific proteomes

To test whether differences in compensation for protein dilution plays a role in additional contexts, we collected *in vivo* pulse labeling samples from four tissues with varying growth rates, Fig. 3a. These samples included the brain, one of the tissues with the lowest reported cellular turnover, as well as the bone marrow, one of the fastest renewing systems in the body^26^. Samples were collected from 10 different mice that span both genders and two different age ranges, 4 and 24 months.

**Figure 3.**
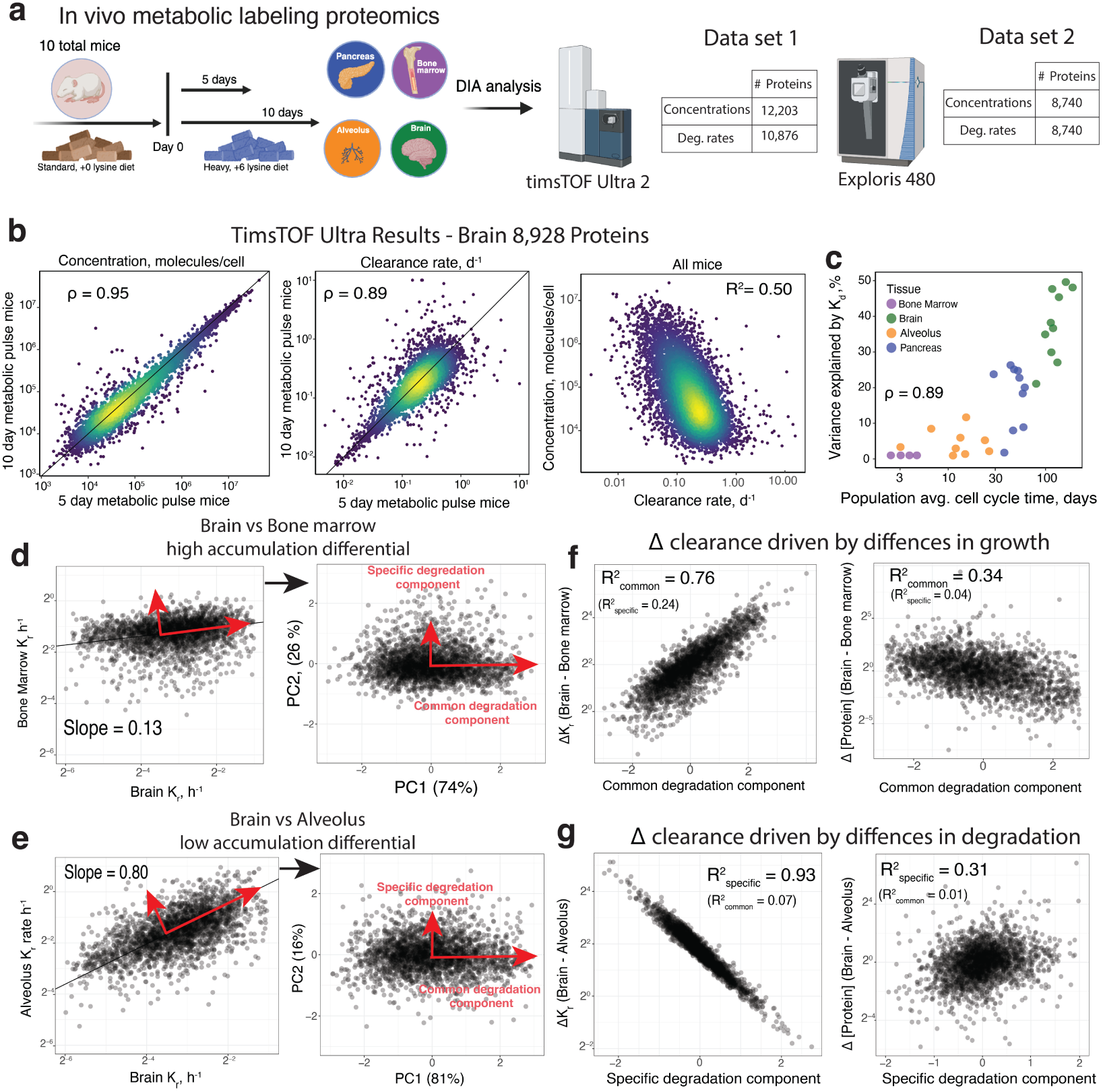
Tissue-specific protein dilution and degradation rates explain protein abundance variation across tissues. **a**, Protein clearance and synthesis rates were measured from murine tissue samples using *in vivo* metabolic pulse labeling with heavy lysine. Samples were then analyzed by (i) timsTOF Ultra 2, which quantified over 12,000 total protein abundances and over 10,000 degradation rates across all tissues and (ii) Exploris 480, which quantified 8,740 abundances and degradation rates. **b**, Reproducibility of protein abundance and degradation rate estimates for 8,929 proteins in the brain. Protein degradation and abundance are inversely related across the proteome, *R*^2^ = 0.50. **c**, Variability in degradation rates and clearance rates for each tissue. Growth rates are reflected by size of points and explain differences between degradation and clearance variability for alveoli and bone marrow. **d**, When plotting clearance rates for brain against bone marrow, the shallow slope indicates reduced variance in bone marrow clearance rates due to high dilution in bone marrow. We then decompose clearance rate trends into a common component reflecting the shared axis of variation in protein degradation rates and a specific component reflecting differences in degradation. **e**, These trends are also shown in comparison between lung and bone marrow where the growth differences are less significant and thus the slope is less shallow. **f**, When growth differences are substantial, the common degradation component drives clearance differences. This in turn contributes to abundance differences between the tissues. **g**, When growth differences are small, the specific degradation component drives clearance and abundance differences between tissues.

In order to draw accurate conclusions from the data, we sought to account for measurement noise. To this end, we estimated abundance and clearance rates from samples digested by either Lys-C only, or both Lys-C and trypsin. Samples digested by different proteases generate different peptides and thus share less systematic bias from factors such as ionization and digestion efficiency compared to technical replicates^27^. Because Lys-C and trypsin share a subset of generated peptides, this will lead to an underestimation of bias when estimating protein abundance. Underestimating this abundance bias should lead to conservative estimates of percentage variance explained by clearance. We also made measurements via data independent acquisition (DIA) MS using two different mass spectrometer instruments and quantification strategies. Trypsin digested samples were analyzed by a timsTOF Ultra 2 mass spectrometer where quantification was performed at the MS2 fragment level while samples digested exclusively by LysC were analyzed on an Exploris 480 orbitrap mass spectrometer utilizing MS1 level precursor ion quantification. Utilizing these two strategies allows us to account for different systematic biases from factors such as precursor ion interference and peptide fragmentation biases that differ between methods. We applied the spearman correction which allowed us to leverage the similarity between these different methods to estimate and correct for measurement noise when estimating percent variation explained by clearance.

Our primary dataset consisted of 34 samples digested with trypsin and analyzed on the timsTOF Ultra, and 21 samples digested with Lys-C and analyzed on the Exploris 480, Table EV 1. As a secondary data set, we also analyzed the same trypsin digested samples acquired on the Exploris 480, Table EV 1. Within each tissue, all samples came from different mice, and no technical replicates were included. Lys-C measurements from the pancreas were excluded because of the large number of tryptic peptides detected, which we attributed to the endogenous trypsin in the tissue. In samples acquired on the timsTOF Ultra, we quantified the abundances of over 12,000 proteins across all samples, with up to 10,000 in individual brain samples, Fig. EV4a,b. We quantified 10,876 protein half-lives across all samples, with up to 7,950 per tissue sample, Fig. EV4c. From the Lys-C and trypsin digests analyzed on the Exploris 480, we quantified abundances and clearance rates for over 8,740 proteins across all samples, with roughly 5,000 per sample, Fig. EV4c.

To understand how tissue type and biological variability shaped protein abundances in each dataset, we first correlated absolute protein abundances between all pairs of samples across the full set of quantified proteins, Fig. EV4d,e. In both the trypsin and Lys-C data, samples clustered clearly by tissue, indicating that, as expected, tissue type is the dominant source of variation. To gauge technical reproducibility in the absence of technical replicates, we randomly split the samples of each tissue into two groups and averaged within each group to reduce within-tissue biological noise. This revealed high reproducibility within every tissue (Pearson *r >* 0.94 for all tissues; Fig. EV4f,g). Agreement of protein fold changes between tissues was strongest between bone marrow and alveolus and most distinct for the brain, as previously observed^28^.

Reproducible measurements are not necessarily accurate, however, since systematic biases can distort protein levels. To assess quantitative accuracy in the presence of biases such as ionization efficiency and precursor interference, which affect different peptides differently, we examined the agreement between peptides mapping to the same protein. For each protein with more than two peptides, we split its peptides into two groups and correlated their relative abundances across tissues, Fig. EV4h,i^29^. We observed strong agreement in both datasets, whereas correlations between peptides mapping to different proteins were on average close to zero. However, additional systematic biases can be incurred based on quantitation method, Ms1 vs Ms2, and MS instrument and digestion enzyme used. For each tissue, we compared protein abundances from digest conditions and observed strong agreement between the estimated protein abundances, in line with expectations from Lys-C and Trypsin proteases^27^, Fig. EV5a. For clearance rate measurements, to further mitigate the influence of noise, we filtered estimates within each digest and tissue for proteins with a coefficient of variation (CV) below 20 %. This set contained roughly 5,000 proteins for each tissue. We then compared the consistency between digests for the intersected subset of low CV observations Fig. EV5b. We observed good consistency for brain and lung measurements, *ρ* = 0.82 and *ρ* = 0.78 respectively. The consistency was notably lowest for bone marrow. This is primarily due to the small dynamic range in clearance rates which are primarily driven by dilution. A comparison of clearance rates across tissues showed smallest amount of agreement between bone marrow and other tissues, Fig. EV5c. This likely reflects the challenges in measuring clearance rates in rapidly dividing cells.

The population average cell growth for each tissue was estimated under the assumption that the histone degradation rates are much smaller than dilution of histones upon cell division^30,31^. Therefore, we defined *k_g_*for each sample as the average clearance rates of histones H3 and H4. Indeed, this assumption is directly supported by recent single-cell metabolic pulse-labeling measurements^30^, where heavy/light histone ratio resolves two discrete populations of cells. Cells which had not divided retained almost exclusively old (light) histones, and those that had divided once, whose histones showed an approximately even heavy/light split. The near-complete retention of light histones in non-dividing cells confirms that histone degradation is negligible over the labeling period, so histone clearance reflects dilution by cell division and provides a direct readout of the growth rate *k_g_*. We observed high consistency between estimates made from each digest condition, Fig. EV5d. The average population doubling times ranged from approximately 1 day for bone marrow to 200 days for some brain samples. Note, that these estimates (and thus the associated assumption) do not affect one of the results about protein clearance.

As shown in Fig. 1, protein abundances are substantially controlled by the rate of clearance in cells with slow growth rates. To evaluate this trend *in vivo*, we first examined brain tissue, which showed the slowest rate of cell division. In the data acquired on the timsTOF Ultra, we observed consistent abundance and degradation rate measurements for 8,928 proteins, Fig. 3b. We again observed a strong relationship between these protein abundances and degradation rates, *ρ* = *−*0.65. When accounting for measurement reliabilities (Fig. 3b), this correlation corresponds to *R*^2^ = 0.5. This estimate suggests that 50 % of protein abundance variation in the brain is controlled by protein degradation.

Because we observed a broad range of doubling times, we next examined how doubling times affected the relationship between degradation and protein abundance. To examine this relationship, we utilized average abundance and degradation measurements in the space of intersected proteins from both digests. To account for measurement noise, we applied the Spearman noise correction, which rescales the variance explained (*R*^2^) by each measurement’s reliability, defined as the Pearson correlation between independent replicate or cross-method (Lys-C vs. trypsin, MS1 vs. MS2) measurements of the same quantity. This correction relies on the additivity of variances of uncorrelated variables and is rigorously correct and precise when the noise is not correlated to the signal^29^. Consistent with our earlier results, clearance rates weakly contribute to protein abundances in rapidly proliferating cells such as the bone marrow. Conversely, clearance contributed up to 50 % of abundance variation in slowly dividing tissues such as the brain. When looking across all tissues, we observed a continuous trend with a strong correlation, *ρ* = 0.89, Fig. 3c. This trend generalizes beyond our four tissues. Applying a similar analysis to a previously published turnover atlas spanning 14 mouse tissues and brain regions^12^, we found that the per-tissue contribution of protein half-life to abundance again increased continuously with doubling time (estimated from histone turnover), Pearson *r* = 0.81, Fig. EV6.

We then set out to determine how these growth rate differences shape clearance and abundance differences across tissues. First, we plotted clearance rates for the brain against the bone marrow, Fig. 3d. We observed that while clearance rates were correlated, *ρ* = 0.40. The slope estimated by total least squares is shallow, 0.13, indicating reduced variability in clearance rates for the bone marrow. This reduced variability is consistent with the faster cell growth in bone marrow, and thus larger contribution of the constant dilution term to the clearance rates.

Despite the shallow slope, the correlation of the clearance rates defines a clear axis of similarity along the line passing through the data points. This axis captures the shared variance in clearance rates, which we termed a common component. The tissue-specific differences in degradation rates are reflected by the variance orthogonal to the common component, and we term it specific component. Mathematically, the common component and specific component correspond to the principal components of the data, PC1 and PC2. The common component reflecting shared clearance variation explains the majority of the variance, 84% on PC1. When we compare the brain to the lung, we see that the slope is less shallow, 0.80, reflecting the smaller impact of dilution for the lung Fig. 3e. Again, the common component explains the majority of the variance in clearance rates. We note that although the common component explains the majority of variance in clearance rates between tissues, it can obscure significant differences in protein degradation that are masked by the differing degrees of dilution between the compared tissues. We interpret this common component as dilution, rather than coordinated changes in degradation machinery, because it corresponds to a near-uniform shift in clearance across all proteins and its magnitude scales with the growth-rate differential between tissues; a contribution from correlated changes in shared degradation pathways that affects all proteins in a similar way cannot be fully excluded.

Although the common component explains the majority of variance in both tissue comparisons, its relationship to clearance and abundance differences between the tissues differs. In tissues with a large growth rate differential such as the brain and bone marrow, the common component explains 76 % of the clearance differences Fig. 3f. As these differences reflected the axis of shared variation in degradation rates, they are directly attributable to dilution differences. Similar to observations made in resting and activated B-cells, we found that the common component explains a substantial fraction of abundance differences between the brain and bone marrow, *R*^2^ = 0.34, Fig. 3f. This again suggests that cells do not fully compensate for the reduction in protein accumulation in fast growth states. However, when tissues have similar clearance rate variability, the specific degradation component explains the majority of clearance rate differences, Fig. 3g. Here, the specific component driven by degradation rate differences explains 20 % of protein abundance differences between the brain and lung Fig. 3g.

Differences in clearance driven by degradation are most pronounced when comparing rates for the two tissues with the lowest growth rates, brain and pancreas. Here, we observed a modest correlation between clearance rates, *ρ* = 0.65, demonstrating significant degradation rate differences, Fig. 4a. Protein fold changes between the pancreas and brain were strongly predicted by clearance differences, *R*^2^ = 0.32, Fig. 4b. This trend was the strongest for any pairwise comparison, Fig. 4c.

**Figure 4.**
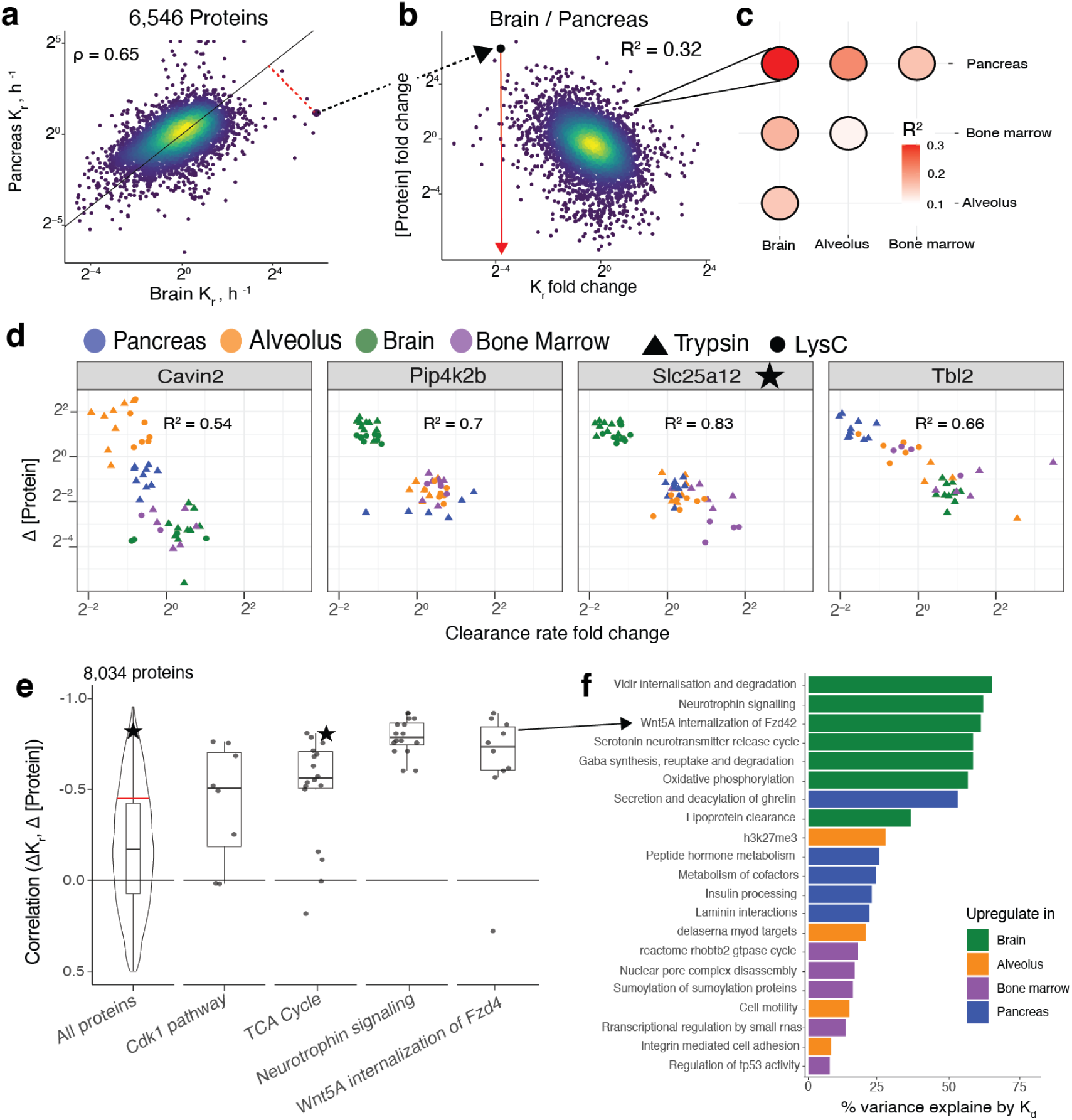
Tissue specific abundances of many proteins are primarily determined by their tissue specific clearance rates. **a**, Protein clearance rates for pancreas and brain plotted against each other show a correlation of 0.65. **b**, Differences in clearance rates between brain and pancreas explain differential protein abundances, *R*^2^ = 0.32. **c**, Similarly, differences in clearance rates explain differences in abundances between other tissue types to varying degrees. **d**, For some proteins, abundance across tissues is almost entirely explained by variation in their clearance rates. **e**, On average, the raw correlation between clearance rates and correlation profiles across protiens analyzed on the timsTOF was 0.20. Accounting for measurement noise via the Spearman correction on the intersect between Lys-C and trypsin data sets estimates the average correlation as 0.46, depicted by the red line. The star depicts the correlation of Slc25a12 shown in panel **d**. This protein participates in the TCA cycle, whose enzymes are primarily regulated by degradation. **f**, GO terms with significant contribution of degradation rates are summarized along with what tissue the pathway proteins are more abundant. The results in this figure are from Dataset 1 (MS2-level quantification, timsTOF Ultra), and results from analogous analysis of Dataset 2 (MS1-level quantification, Exploris 480) as are shown in Fig. 5 as an independent replicate.

We next sought to examine the extent to which variations in the abundances of individual proteins across samples could be explained by degradation variation. We identified many proteins whose abundance could be mostly explained by clearance, *R*^2^ *>* 0.70, Fig. 4d, Table EV2. Because we measured more proteins in samples from the timsTOF Ultra, we did not correct the correlations for individual proteins for measurement noise. When examining the distribution across all proteins, we found an average correlation of 0.20, Fig. 4e. We used the spearman correction factor to estimate a noise-corrected mean correlation in the space of proteins measured in both the timsTOF trypsin data set and the Lys-C Exploris 480 data set, resulting in a correlation value of 0.46 (see methods). To elucidate this trend, it was important to control for average differences in clearance rates. Clearance rates for all proteins will be higher in bone marrow compared to brain, but it is the differences relative to the average clearance that will impart specificity to protein abundances.

Additionally, we observed that protein variation for several pathways is substantially controlled by protein degradation. Increased abundance in the brain of proteins from pathways such as Neurotrophilin signaling and WNT5A signaling can be almost entirely explained by their slower degradation rates, Fig. 4f. Several other pathways such as serotonin and insulin receptor recycling are also up regulated in the brain. Other pathways with significant control by degradation such as Insulin processing and peptide hormone metabolism were up-regulated in the pancreas. Overall we identified 141 pathways whose abundance is controlled by degradation at more than 20%, Table EV2. A replicate analysis with additional samples digested by trypsin and collected on the Exploris 480, Table EV1, shows similar results (Fig. 5) on a shallower subset of proteins.

**Figure 5.**
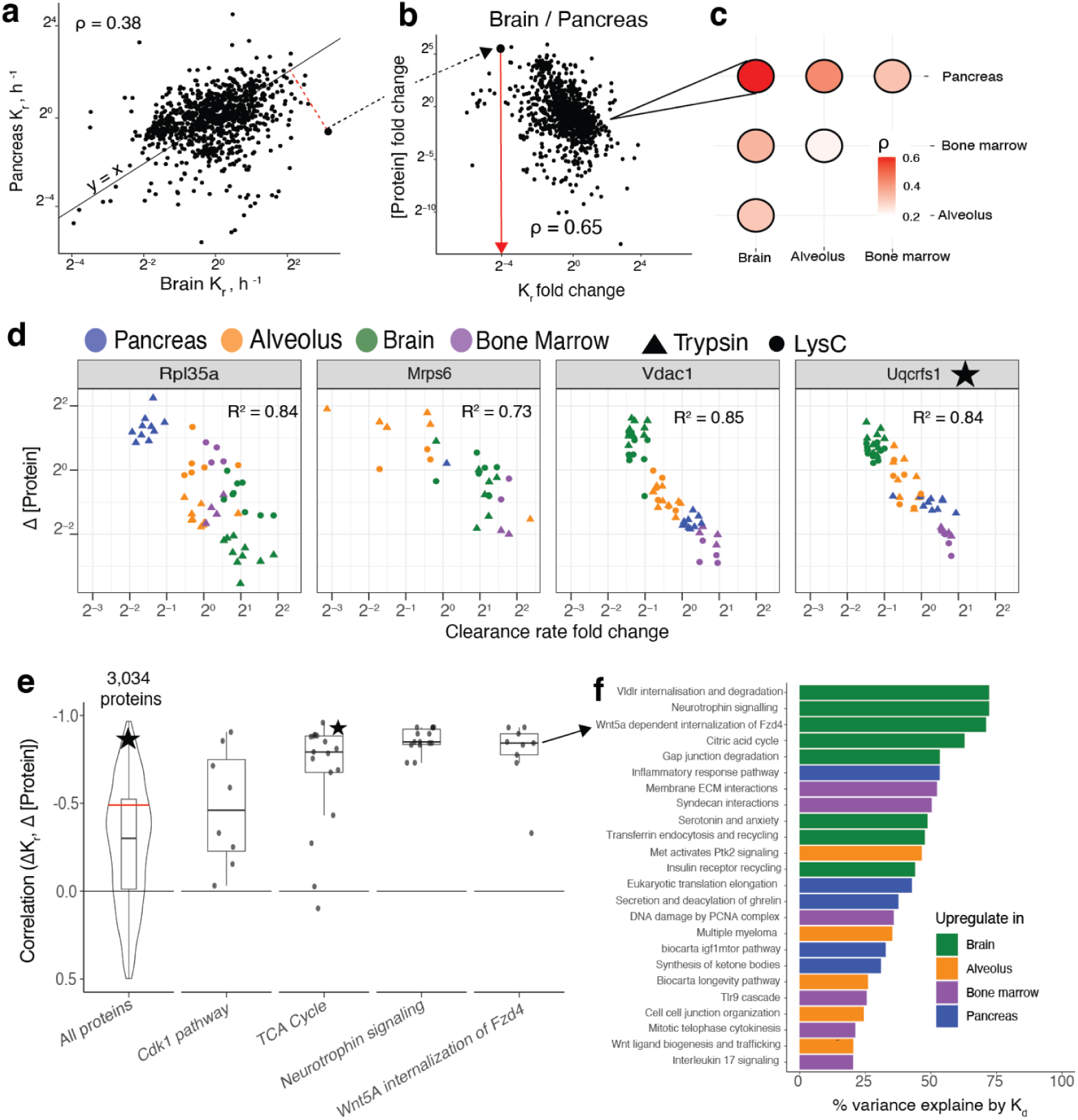
Tissue specific abundances of many proteins are primarily determined by their tissue specific clearance rates. The results in this figure are analogous to those shown in Fig. 4 but are based only on Dataset 2 (acquired by Exploris 480) and included as an independent replicate. **a**, Protein clearance rates for pancreas and brain plotted against each other show modest correlation. **b**, Differences in clearance rates between brain and pancreas explain differential protein abundances, Pearson correlation *ρ* = 0.65. **c**, Similarly, differences in clearance rates explain differences in abundances between other tissue types to varying degrees. **d**, For some proteins, abundance across tissues is almost entirely explained by variation in their clearance rates. **e**, On average, the correlation between clearance rates and correlation profiles is 0.35. When correcting the correlation for measurement for consistency of Lys-C and trypsin digests via the spearman correction, the average is 0.50 depicted by the red line. The star depicts the correlation of Uqcrfs1 shown in panel **d**. This protein participates in the TCA cycle, whose enzymes are primarily regulated by clearance. **f**, GO terms with significant contribution of clearance rates are summarized along with what tissue the pathway proteins are more abundant.

### Proteoform group specific degradation rates influence proteoform group abundances

Having found that degradation rates substantially influence abundance across different proteins and samples, we sought to further investigate this influence across proteoforms encoded by the same gene. Such analysis is challenging due to the high sequence homology of proteoforms, but our high protein sequence coverage, quantifying on average 10 lysine containing peptides per protein, makes it feasible, Fig. 3. We focused on peptides mapping to proteoform groups generated by alternative splicing in the brain. This choice reflects the prominent role of alternative splicing (AS)^32^ and the pronounced role of degradation in shaping protein abundances in the brain, Fig. 3b.

To determine a set of relevant AS events, we utilized the VastDB database to curate a set of specific AS events for the brain^33^. We selected all events with at least 20 % difference in percent spliced in (PSI) values compared to the all other tissues. For these events, we mapped quantified peptides to potential splice isoforms. Most peptides mapped to multiple AS proteoform, though some peptides mapped uniquely to a single proteoform, Fig. 6a. Peptides were then sorted into groups based on their shared set of potential isoforms. Testing differences in degradation rates between the groups for 500 different alternative splicing events yielded 172 events (34%) containing proteoforms with significantly different degradation rates, FDR *<* 0.05, Table EV3.

**Figure 6.**
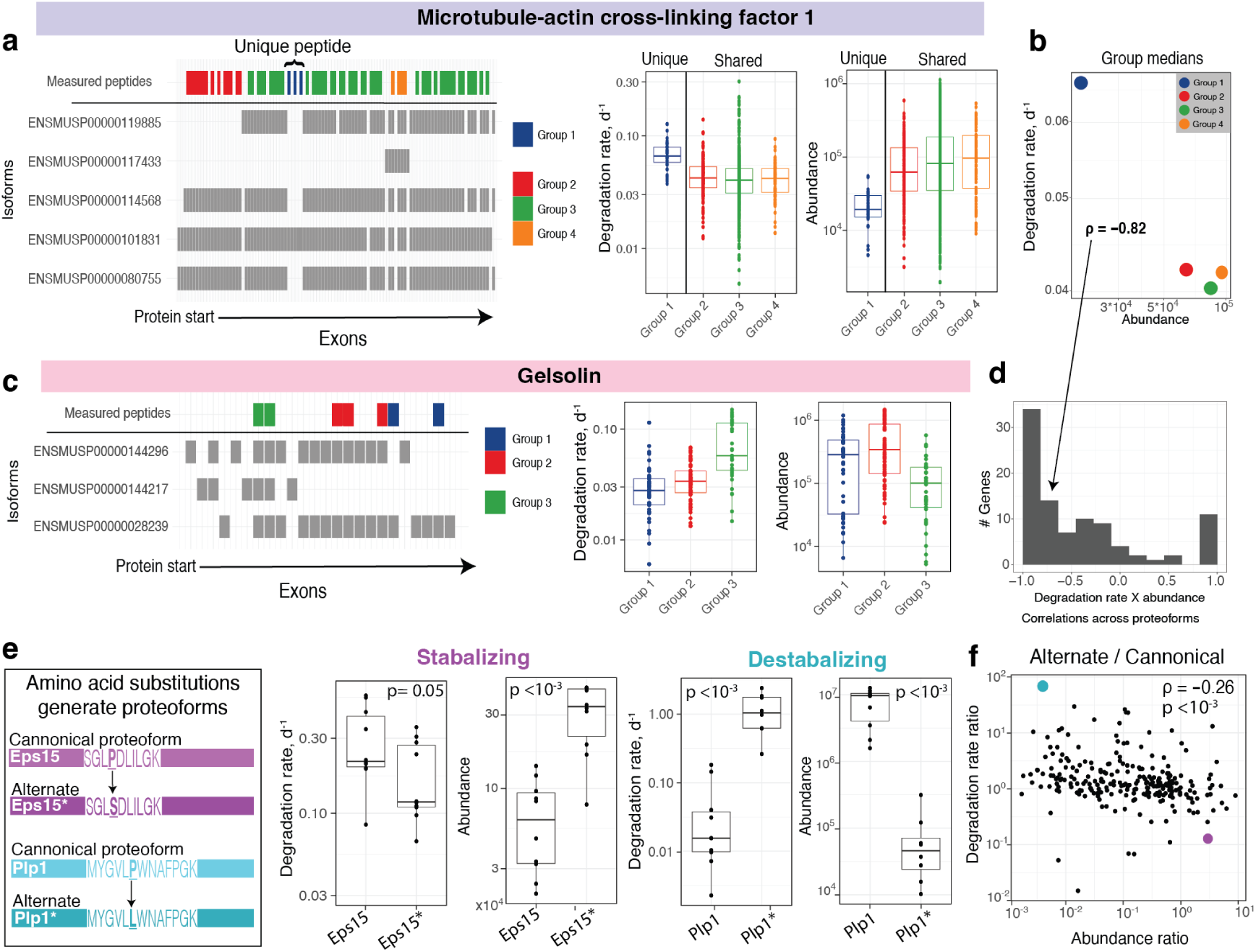
Proteoform-specific degradation rates influence abundance across proteoforms. **a**, Measured peptides mapping to different proteoforms of the gene Microtubule-actin cross-linking factor 1 are plotted by the position of the exon encoding their sequence. Colors correspond to potential set of proteoforms to which the peptide could belong. Group one peptides are unique to proteoform ENSMUP00000101831. Group 1 peptides have higher degradation rates than peptides from other proteoforms and correspondingly have lower abundance. **b**, Correlation between the average degradation rates and abundances for peptides from the 4 proteoform groups, *ρ* = *−*0.82. **c**, Peptides mapping to different groups of proteoforms from the gene Gelsolin show varied degradation rates. Again the group of peptides with the fastest degradation rate has correspondingly low abundance. **d**, Correlations between the median degradation rate and abundance for peptides mapping to a shared potential set of proteoforms for the 172 events with significant differences in degradation across proteoforms. A strong negative trend, median *ρ* = *−*0.76, suggests proteoform abundances from a share genes are controlled by degradation. **e**, Alternate translation of Eps15 and Plp1 results in proteoforms with different amino acids. These single amino acid differences stabilize the alternate proteoform of Eps15 leading to higher abundance while destabilizing the alternate proteoform of Plp1, leading to reduce abundance. **f**, The differences in degradation rates and in abundances across all pairs of alternate and canonical proteoforms are inversely correlated, *ρ* = *−*0.25.

Peptides mapping to the isoform ENSMUSO000000112568 from Microtubule-actin crosslinking factor 1 showed higher degradation rate compared to peptides mapping to different splice isoforms, Fig. 6a. Additionally, the abundance of these peptides was significantly lower, p-value *<* 10*^−^*^4^, suggesting that proteoform abundance is influenced by proteoform-specific degradation. We visualized this influence by the medians of the degradation rates plotted versus the medians of abundances for each group, Fig. 6b, and quantified it but the correlation, *ρ* = *−*0.82. Another example from examining proteoforms of Gelsolin shows three groups of peptides mapping to different potential subsets of proteoforms Fig. 6c. Only peptides from group 3 map to ENSMUSP0000144217. These peptides show faster degradation rates and correspondingly lower abundances. Comparing abundances and degradation rates of peptides mapping to different proteoforms across all 172 significant events showed a median correlation of *ρ* = *−*0.76, Fig. 6d. Together, these results suggest that the stability of the different splice isoforms inside the cell may be an important factor in controlling their abundance in addition to variation in splicing frequencies. However, we acknowledge that without matched, isoform-resolved RNA-seq we cannot confirm which isoforms are present, nor fully separate the contribution of differential splicing from differential clearance to the observed proteoform-group abundances. Our data do establish that clearance rates differ significantly between peptide groups and covary inversely with their abundances, but quantifying the relative contributions of splicing and clearance will require matched transcript-isoform measurements^34^.

In addition proteoforms generated from alternative splicing events, we sought to analyze proteoforms arising from alternate translation, manifested as single amino acid substitutions. These proteoforms correspond to incorporating amino acids deviating from the genetic code as previously reported^14,35^. We used deep DDA data from murine tissues^33^ to identify protein products of alternate RNA decoding, which are detected as single amino acid differences from the sequences predicted by in silico translating mRNAs with the genetic code; we used the approach previously reported by Tsour et al.^14^ to identify and validate them, and then quantified their abundance and degradation rates using our timsTOF Ultra dataset. The existence of such proteins resulting from alternate RNA decoding has been reported recently and not well established, but we nonetheless proceeded to investigate the potential role of protein stability in their abundance.

This approach led to the detection and quantification of 306 unique peptide sequences with single amino acid substitutions in addition to their canonical counterparts. Most quantified substitutions, such as the *P* ↦ *S* substitution in Eps15, stabilized the alternate proteoform, leading to increased abundance compared to the canonical form, Fig. 6e. Other substitutions such as a change of proline to leucine in Plp1 destabilized the resulting proteoform, partially contributing to a decrease in abundance compared to the canonical form. To extend this analysis systematically to all proteoforms, we first derive an estimate for the abundance of the alternate relative to the canonical proteoform by computing their ratio, termed the ratio of amino acid substitution (RAAS)^14^. The distribution of alternative sequence intensities ranged from 1/1000th to 10 times the intensity of the canonical sequence. These abundance differences correlated *ρ* = *−*0.26 significantly to the corresponding differences in degradation rates between proteoforms. This again suggests that proteoform-specific degradation rates contribute to determining proteoform abundance.

## Discussion

We measured and analyzed protein abundance and degradation rates at the cell type level and at the whole tissue level across protein products encoded by about 10,000 genes. The results demonstrate: (i) Growth-rate dependent impact of protein clearance on the abundance variation across the proteome, (ii) At slow growth, this impact may account for up to 50% of the measured absolute protein abundance variation across the proteome, much greater than previously appreciated, (iii) Differences in clearance rate caused by differences in degradation or secretion and cell growth substantially contribute to variation in relative protein levels across tissues, and (iv) Protein degradation significantly impacts the abundance of proteoforms from the same gene, thus providing evidence for a molecular mechanism that can explain why proteins are detected from only a few AS isoforms^36^. To enhance robustness, we acquired datasets using multiple proteases and MS instruments. The comparison of these datasets, using MS1- or MS2-level quantification indicated remarkable reproducibility of our results and interpretations as shown in Fig. 4 and Fig. 5.

Our analysis and data indicate that protein degradation impacts protein abundance comparably to protein synthesis at slow growth. This impact stems from the fact that proteins accumulate inversely proportional to their rate of degradation. In growing cells, however, protein accumulation is reduced as dilution becomes a major source of protein clearance. To prevent the reduction in accumulation from altering protein abundances at fast growth, synthesis rates must compensate and increase proportionally to the reduction in accumulation. In proteins like histones, such compensation almost fully offsets the reduction in accumulation, Fig. 2c,d. For other proteins we observe only partial compensation, sharply contrasting with previous reports emphasizing the role of compensation^4,5^. This partial compensation does not offset the majority of changes in clearance rates caused by increased dilution. Our measurements and analysis quantify this effect for the first time for thousands of proteins in physiologically relevant conditions. Thus, we establish the role of clearance differences caused by dilution in shaping cell type and tissue specific proteomes.

We find that this phenomenon explains a substantial amount of variation in protein abundances. It contributes 37% of abundance change between resting and activated B-cells (Fig. 2) and 34 % of the abundance changes between brain and bone marrow tissues (Fig. 3). This raises the interesting possibility that long lived proteins are generally less needed or inhibitory for high growth states and more important for low growth states. While it has been posited that certain proteins such as those involved in signaling require high turnover to allow cells to adapt quickly to environmental stimulus^6^, our observation provides potential explanation in the opposite trend. Proteins crucial for slow growth such as the proteins in the electron transport chain may have preferentially evolved to have longer half lives. While our analysis focused on healthy tissues, it has clear implications for diseases: Decease in degradation rates can substantially increase proteins abundance, as observed in Alzheimer’s disease^37,38^.

We also report large and context-dependent roles of protein degradation both across cellular states, across different proteins, and across proteoforms. While the possibility for such extensive regulation has been suggested by protein variation that is unexplained by mRNA variation^29,39–41^, our analysis focused on explaining protein variation by measured protein degradation rates. The results emphasize a larger than previously described role for protein degradation in determining protein abundance variation. They also explain the smaller roles previously reported in rapidly growing and dividing cells^1^ or cells growing in size^3^. Our results did not include shrinking cells, such as erythroblasts transforming into erythrocytes^42^, but our results predict that protein degradation is likely to play an even more dominant role. Our data demonstrate that variation in protein degradation rates is an important global mechanism for shaping protein and proteoform abundances, Fig. 4 and Fig. 6. This large contribution helps explain the high abundance of stable proteins resulting from alternate RNA decoding^14^. Such analysis of proteoforms is challenging but greatly aided by the large number of peptides quantified for many proteins in our dataset. Future analysis can benefit from the recently published framework utilizing full length RNA sequencing and proteoform isolation^34^. Together, our results motivate future work on understanding the biological impacts of protein degradation across diverse contexts.

Our tissue analysis faces limitations, such as averaging across the diverse cells making up the analyzed tissues. For example, tissues like the brain comprise non-proliferative differentiated cells and proliferative cells. This complicates interpretations as it is challenging to disentangle the exact amount of dilution each quantified protein faces. Single-cell measurements of degradation, growth, and abundance rates in vivo add additional context to understanding when synthesis rates must compensate for protein dilution^43^. Advances in the single-cell measurement technology and data modeling approaches are likely to significantly expand the emerging principles of protein abundance regulation^44^.

## Data availability

All raw data and search engine outputs can be found on MassIVE with ID: [MSV000097050](https://massive.ucsd.edu/ProteoSAFe/dataset.jsp?task=732135590e454112b315f6e610891080). EV tables and all additional processed data needed for reproducing the analysis can be found on Zenodo under DOI https://doi.org/10.5281/zenodo.21444972.

## Code availability

The code used for data analysis and figures is at: github.com/Slavovlab/Protein-Clearance

## Acknowledgments

We thank Yanxin Xu and Bin Zhang for help with mouse tissue preparation, Zhixun Dou for help procuring mice, Eunice Koo for help with identification, and Alexander Franks for advice on statistical analysis. The work was funded by a Bits to Bytes award from MLSC to N.S., an NIGMS award R01GM144967 to N.S., and a MIRA award from the NIGMS of the NIH (R35GM148218) to N.S, a UH3CA268117 award from NIH to N.S., and R01AG092460 award from NIH to N.S.

## Author Contributions

**Experimental design**: A.L. and N.S.

**Sample preparation**: A.L., S.Z. and P.S.

**Raising funding & supervision**: N.S.

**Data analysis**: A.L. and N.S.

**Writing & editing**: A.L. and N.S.

## Methods

### Mouse model and handling

All mice experiments were performed in compliance with the Institutional Animal Care and Use Committee at Massachusetts General Hospital. Both male and female C57BL/6 mice, either 24-month-old or 4-month-old, were ordered from the NIA. Mice were euthanized with CO2 followed by cervical dislocation. The mouse used was male. Tissues were harvested post-euthanasia and perfusion with PBS.

### Metabolic pulse labeling and tissue collection

Mouse Express L-LYSINE [13C6, 99%] MOUSE FEED kit was purchased from Cambridge Isotopes Laboratories. Mice were first acclimated to the Lys+0 feed for a period of 10 days before switching to the Lys+6 diet for either 5 or 10 days. Mice were fed 5g per day for either 5 days or 10 days before euthanisation. Whole tissues including the pancreas, brain, aveolus, and bone marrow were harvested a total of 10 mice. Descriptions of the age, gender and feeding time for each mice can be found in the meta data file on Zenodo. After harvesting, tissues were stored at −80 until prepared for LC-MS/MS analysis.

### Sample preparation for MS

Samples were prepared based on an optimized SP3 protocol^45^. Briefly, whole tissues were thawed from −80 degree freezer at room temperature in 1 mL of 1x PBS. Samples were then ground using 1 mm diameter glass beads and a tissue homogenizer (BioSpec Mini-Beadbeater). Samples were then brought to 1% SDS and 1x benzonase to break apart DNA. Samples were then brought to 5% SDS and boiled for 10 minutes. 10 micrograms of protein from each sample were then added to SP3 beads for clean up and overnight digestion at 37 degrees C in 20 ng/*µ*l LysC. LysC digest was then split into one fraction for LysC analysis and another fraction for a second overnight digest with Trypsin at 37 degrees C in 10 ng/*µ*l Promega Trypsin Gold. Samples were then dried down and resuspended at a concentration of 750 ng/*µ*l in 0.1 % formic acid for injection to the LC-MS/MS.

### LC-MS/MS data acquisition

All samples were were from both Exploris 480 and timsTOF Ultra separated via online nLC on a Vanquish Neo UHPLC using a 25cm x 75 *µl* IonOpticks Aurora Series UHPLC column (AUR2-25075C18A). Samples were resuspended at a concentration of 500 ng/*µ*l and 1 *µ*l volumes were injected out of glass inserts (ThermoFisher 60180-509) sealed by rubber caps (ThermoFisher C5000-51B).

All samples from both Exploris 480 and timsTOF Ultra used a 95 minute total length method (53 minutes active gradients). 0.1 % formic acid in water was used for Buffer A and a 20% solution of 0.1% formic acid mixed in 80 % acetonitrile was used for buffer B. A constant flow rate of 200nl/min was used throughout sample loading and separation. Samples were loaded onto the column for 20 minutes at 1% B buffer, then ramped to 5 B buffer over two minutes. The active gradient then ramped from 5% B buffer to 25% B buffer over 53 minutes. The gradient was then ramped to 95% B buffer over 2 minutes and stayed at that level for 3 minutes to remove excess peptide and protein residue from column. The gradient then dropped to 1% B buffer over 0.1 minutes and stayed at that level for 4.9 minutes for the re-equilibration.

Samples were analyzed by either a Thermo Scientific Exploris 480 mass spectrometer or the timsTOF Ultra. For the Exploris 480, electrospray voltage was set to 1.8 V, applied at the end of the analytical column. The temperature of ion transfer tube was 250 *^o^C* and the S-lens RF level was set to 80. MS1 scans had the following parameters: 140k resolution, 3e6 AGC target, and a scan range from 450 to 1258Th. Two sets of DIA windows were used: 21 20Th-wide windows (spanning the space from 450Th to 860Th) and 8 50Th-wide windows (spanning the space from 859.5Th to 1256Th). The DIA windows had the following characteristics: 35k resolution, AGC target of 5e5, maximum injection time determined automatically, fixed first mass of 200Th, NCE of 27, and a default charge state of 2. DIA windows spanned the space from 450Th to 1256Th and included a 0.5Th overlap. For the timsTOF Ultra, the ion mobility range was set from 0.75 *V ∗ s/cm*^3^ to 1.3 *V ∗ s/cm*^3^. The ramp time and accumulation time were set to 50 ms with a ramp rate of 17.5 Hz. 20 MS/MS ramps were used across 60 MS/MS windows covering 350-1250 m/z.

### Interpreting raw mass spectra

To reanalyze of data dependent acquisition MS data from Mathieson et al.^10^, proteomics data from SILAC-labeled human cell lines was downloaded from the publication data repository (B cells: PXD008511, hepatocytes: PXD008512, monocytes: PXD008513, NK cells: PXD008515). Samples were searched by MSFragger via the SILAC3 workflow and MSBooster workflow^46^ against the human protein sequence fasta downloaded from Uniprot in September 2024.

Label free DIA runs of cell cycle fractions were searched with DIA-NN v1.9.0^47^ using an in silico fasta generated library enabled by deep learning. Raw files were searched together with match between runs (MBR). DIA-NN search settings: Library Generation was set to “IDs, RT, & IM Profiling”, Quantification Strategy was set to “Peak height”, scan window = 1, Mass accuracy = 10 ppm, and MS1 accuracy = 5 ppm, “Remove likely interferences”, “Use isotopologues”, and “MBR” were enabled. SILAC was set as a fixed modification on lysine residues using the plexDIA module, and two channels were defined, one at 0 da and one at 6.051984 da. Search parameters were set to 1% precursor FDR and data was then filtered to 1% protein FDR via the output quantity Lib PG Q Value less than 0.01.

We also performed a variable modification search in DIA-NN with the variable SILAC modification specified at 6.051984 da to identify partially labeled peptides that contained multiple lysines due to a missed cleavage. All other settings were the same other than the use of the fixed SILAC modification and channel command.

### Downstream data analysis

#### Analysis of in vitro external data sets

Protein clearance rate data for fibriblasts, B-cells, NK cells, Monocytes, and Hepatocytes was downloaded from supplemental files of Schwanhausser et al.^1^ and Mathieson et al.^10^. Estimates of protein clearance rate are proportional to the ratio of heavy over light peptide intensities, and thus accurate estimation of protein clearance do not depend on accurate knowledge of cell growth rate. For fibroblasts, protein copy numbers were also downloaded from the supplement of Schwanhausser et al. For the other cell types, we computed protein intensity from the fragpipe searched data by first summing the intensities of light and heavy SILAC precursors. We then estimated protein abundance as the median of the top 3 most abundant precursors for a protein. Random noise in the measurements was accounted for using the Spearman correction and the reliability was estimated by correlating replicate measurements as previously described^29^.

To account for sources of systematic bias such as ionization efficiency digestion efficiency in the *in vivo* metabolic pulse experiments, we collected data digested by different enzymes and run on different mass spectrometer instruments. Averaging measurements with different biases allows for averaging out systematic biases. Additionally, the agreement between different methods allows for more accurate estimation of measurement noise, as these estimates are less likely to be falsely inflated by shared biases. This potentially allows for a more significant noise correction. Note that for pancreas samples, because the LysC digestion contained many tryptic peptides, we used reproducibility of replicate trypsin samples in comparisons including pancreas. This will likely lead to underestimation of errors and thus underestimation of the percent variance explained by degradation.

#### Estimating clearance and cell division rates in vivo

To estimate the change in the fraction of light lysine in the unbound lysine pool over time, *γ*(*t*), we first calculated the fraction of heavy lysine in the free pool at our two time points. To do this, we utilized the average ratio of partially labeled to fully labeled peptides as done by Jovanovic et al.^3^. We then fit an equation of the form *γ*(*t*) = 1 *−* 0.5*e^−at^ −* 0.5*e^−bt^* to estimate the change in available light lysine over time as previously developed and used in ref.^11^. The equation for amino acid availability was fit separately within each tissue type and within each age group to account for tissue and age specific amino acid incorporation^48^.

Having solved for the change in the free lysine pool over time, we then formulated the change in any given light peptide over time 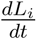 via the equation:

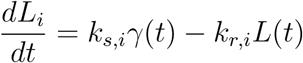

We then integrate this equation to solve for *L_i_*(*t*) for the initial condition *L*(0) = *L*_0_ where *L*_0_ is defined as the sum of heavy and light intensities under the steady state assumption that overall protein concentration is not changing over time. Integrating and further substituting *k_s,i_* via the steady state equality 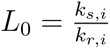 This gives the following equation:

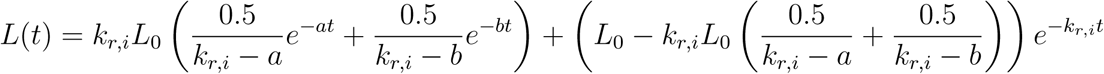

The arithmetic was performed in Mathematica to obviate manual integration errors. Lastly, this equation was then solved for via a non-linear solver to find *k_d,i_* for each peptide from each sample.

#### Math for model of clearance upon growth

For the following derivations, are more comprehensively derived in Baum *et al.*^5^. Briefly, in cells that are not growing, under the steady state assumption protein concentrations can be formulated as a balance of rate of synthesis and degradation.

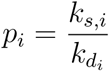

In cells that are growing, the steady state model can be maintained under the assumption that protein concentrations remain constant as cells double the amount of all proteins at an equal rate. However, under this regime, we need to introduce an additional term to reflect protein clearance due not only to degradation, but to dilution as well.

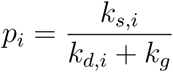

The dilution rate *k_g_* is constant for all proteins and is inversely proportional to the rate of cell division, *T_CC_*, *k_g_* = *log*(2)*/T_CC_*. For both models, we define the overall amount of protein clearance from degradation and dilution is 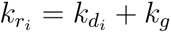 with *k_g_* = 0 in the absence of growth.

When comparing a rapid change from slow to high growth, we have to define the change in both clearance and synthesis rates that contribute to protein concentrations. The difference in clearance rates for all proteins Δ*K_r_* is defined as:

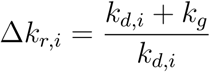

This assumes no significant changes in degradation rates as a cell increases its growth rate. While this assumption may not hold, because the influence of degradation is diminished in the fast growth regimes, the ability of degradation rate changes to compensate for dilution is limited.

The change in synthesis rate has two components. First, there is the overall increase in synthesis that is applied to all proteins, *δ* uniformly to account for the increase in average clearance rate from dilution. Secondly, there is the protein specific change in synthesis, *µ_i_*, that accounts for deviations from the scalar increase, and may compensate for the decreased accumulation of long lived proteins. The final equation describing the difference in clearance

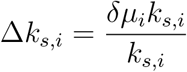

#### Relative contributions of synthesis and degradation to protein levels

To determine the relative contributions of synthesis rates (*K_S_*) and degradation rates (*K_d_*) to protein levels (*P*), we log-transformed the equation 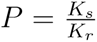 to obtain log(*P*), log(*K_s_*) - log(*K_d_*). The variables log(*K_s_*), log(*K_r_*), and log(*P*) were then mean-centered to remove any intercepts and isolate the variability. We then calculated the relative contribution of either term as the sum of squared residuals from a leave-one-out regression. Further, we corrected *R*^2^ values for the reliability of the data using the Spearman correction as previously described^29^. Briefly, the fraction of variance explained was divided by the reliabilities of each measurement, concentration and degradation rate. The reliabilities were estimated as the correlations between different measurements, such as estimates derived from biological replicates. In the case of the mouse tissues, digestions by lysC and trypsin were also used for reliability estimation.

### Simulation Description

The simulation of our hypothesized protein regulation model from Fig. 1 is described in the Supplemental Information.

### Pathway enrichment analysis

Pathway enrichment analysis was performed on both protein fold changes, regulated protein synthesis changes, and correlations between concentrations and clearance rates. In all cases, a t-test was performed between values from a particular KEGG gene ontology pathway and the null distribution of comparisons for all other proteins. Comparisons were only made if 5 or more proteins were present in a pathway. The distribution of p-values generated from the comparisons was then corrected for multiple hypothesis testing via the Benjamini-Hochberg false discovery rate (FDR) correction. Pathways below 1 % FDR were considered significant.

### Splice isoform analysis

To curate a list of splicing events relevant in the brain, we downloaded a list of PSI by tissue for each gene from VastDB^32^. We filtered for genes with PSI values with a larger than 20 % difference from the average of all other tissues. We then download a list of alternative splicing events for each gene from VastDB, which contained the prominent splice isoforms for each gene. We then filtered through each event to see if peptides from our data mapping to at least two different groups of isoforms were present. For these cases, we tested the distribution of degradation rates for peptides mapping to different subsets of potential isoforms via anova. For the 500 splicing events that were tested, we then adjusted the distribution of p-values with the Benjamini-Hochberg correction and filtered the results at 1% FDR.

### Amino acid substitution analysis

Because our data was collected using data independent acquisition, this posed challenges for denovo identification of alternative translation events as described in Tsour et al., which benefited from existing tools only applicable to data dependent acquisition^49,50^. Thus, to identify these alternative sequences, we curated a list of potential candidates identified from two sources and added these peptides to our spectral library used to analyze our DIA data. First, we took the 456 alternative translation events identified in murine tissues from Tsour et al.^49^. Secondly, we analyzed offline fractionated data dependent acquisition proteomics data generated from murine brain samples using the dependent peptide search and analysis pipeline from Tsour et al.^33^. Briefly, we applied the dependent peptide search and filtered for results corresponding to mass shifts that could correspond to potential amino acid substitutions and were not consistent with any known modifications.

## Expanded View Figures

**Fig. EV1.**
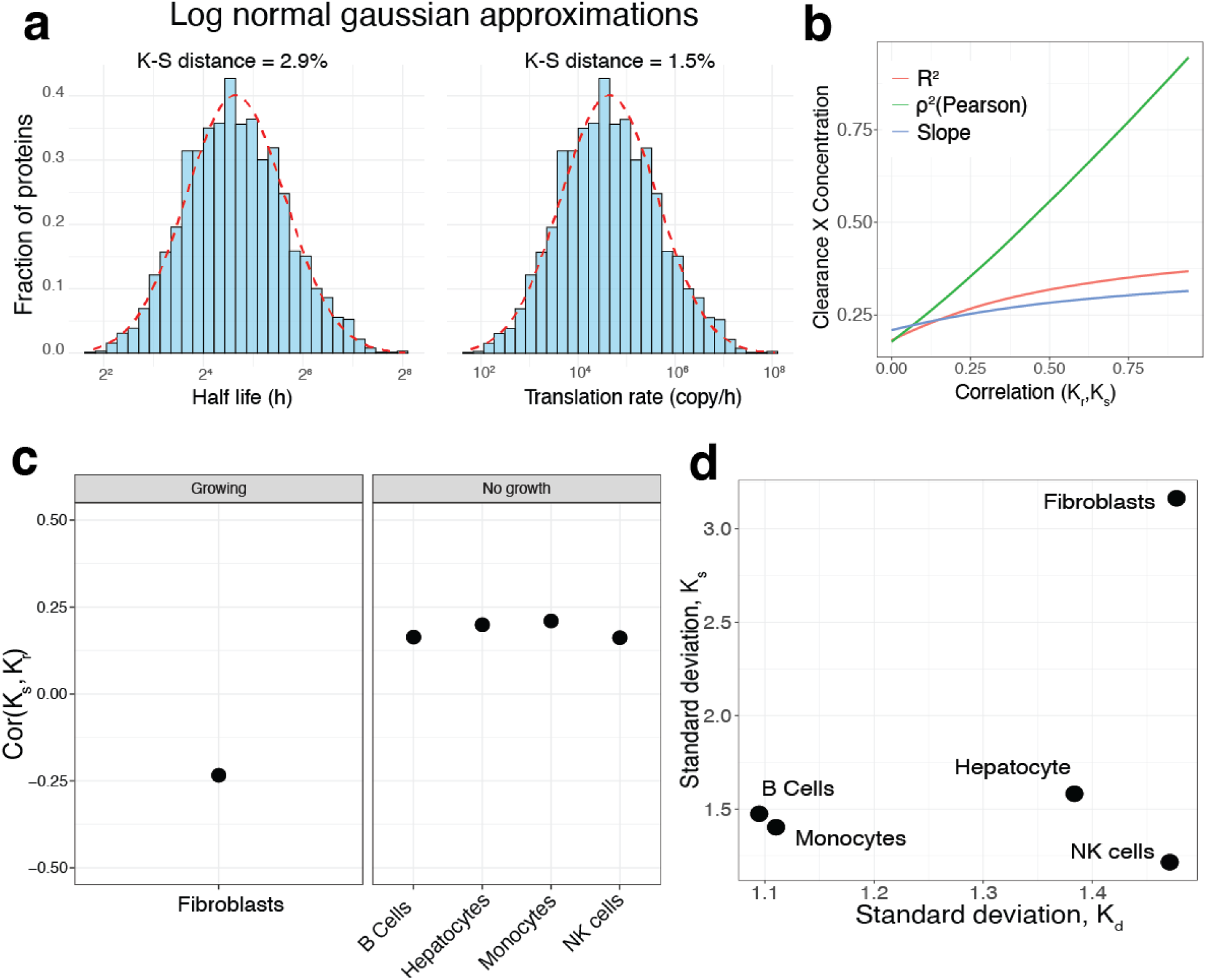
Evaluating characteristics of degradation and synthesis distributions. (**a**), The K-S distances of an approximated log normal fit to the half life and synthesis rate distributions show close concordance differing at most by 3%. (**b**) The *R*^2^ computed as the sum of squared residuals from the relationship *−log*(*K_r_*) = *log*(*P*) does not overestimate the percent variance with increased correlation between growth and synthesis. This can be observed as the slope computed between *−log*(*K_r_*) and *log*(*P*) remains modest and does not approach 1. (**c**) Correlation between synthesis and degradation rates plotted for each cell type. (**d**) Standard deviations of the *log*(*K_r_*) and *log*(*K_s_*) distributions for each cell type.

**Fig. EV2.**
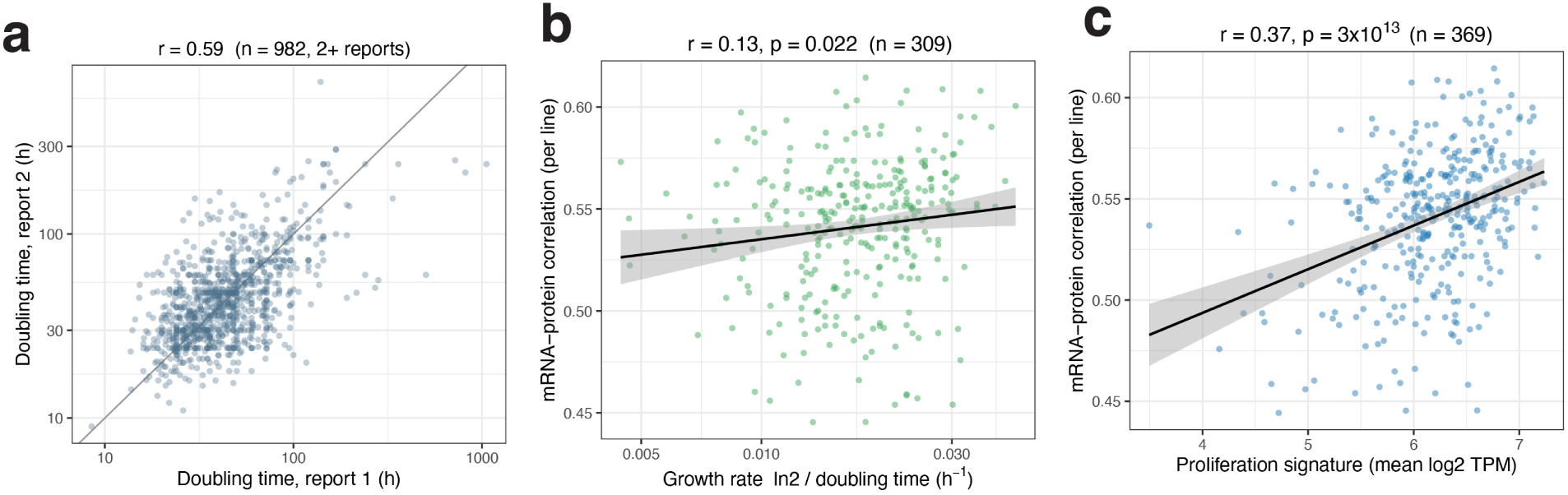
mRNA–protein agreement versus growth rate across cancer cell lines. (**a**) Reproducibility of literature doubling times: two independently reported doubling times for the same cancer cell line, for *n* = 982 lines with at least two reports^18^. (**b**) Per cell line mRNA-protein correlation versus growth rate (ln 2*/*doubling time, from literature doubling times); *n* = 309 lines. (**c**) Per-cell-line mRNA–protein correlation versus a transcriptome proliferation signature (mean log_2_ TPM of proliferation/cell-cycle genes); *n* = 369 lines.

**Fig. EV3.**
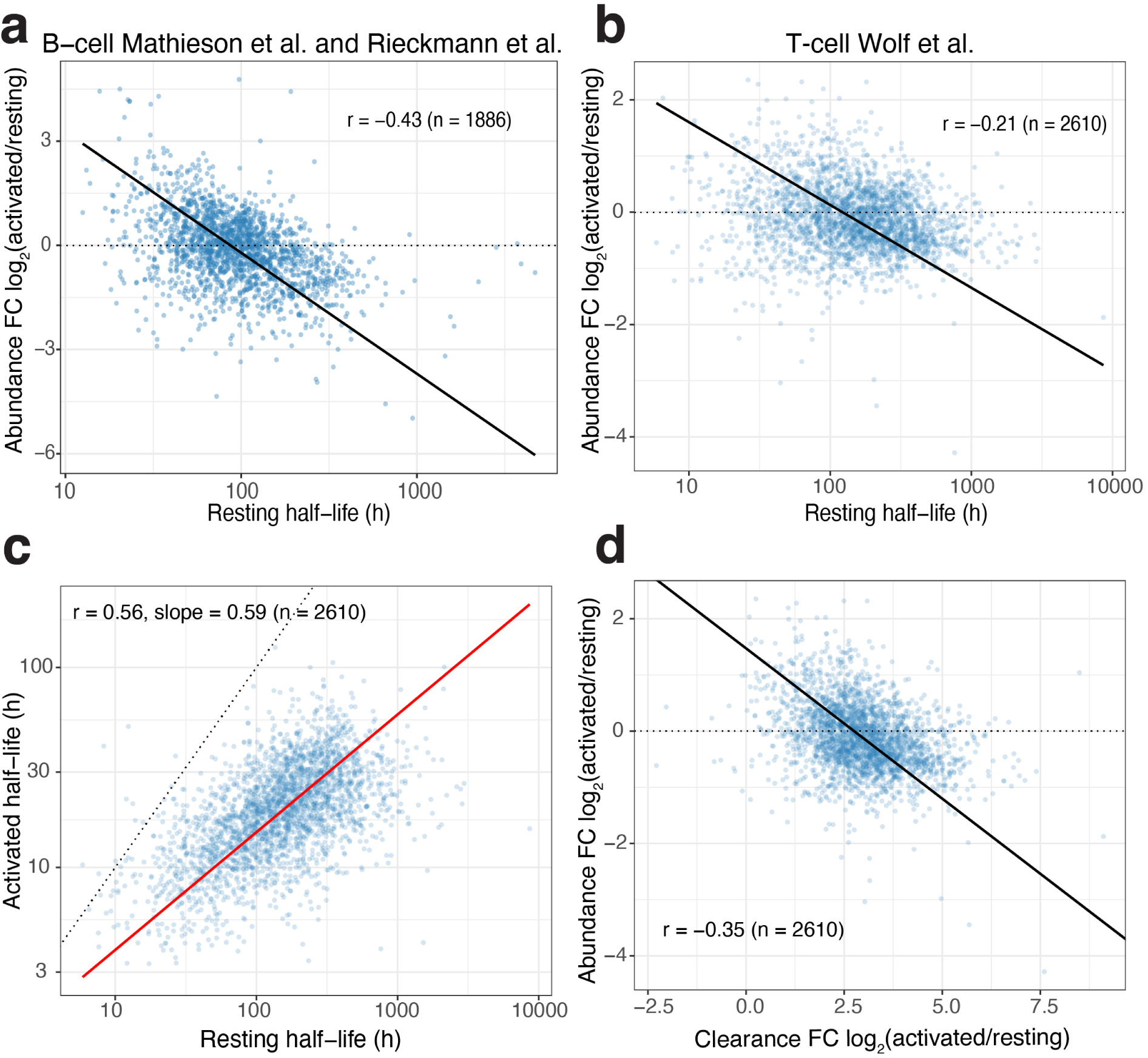
Protein stability and abundance change upon lymphocyte activation. (**a**) Change in protein abundance upon B-cell activation, log_2_(activated/resting), versus resting protein half-life. Resting half-lives are from Mathieson *et al.*^10^ and abundances from Rieckmann^21^ *et al.*; *n* = 1,886 proteins. (**b**) As in (**a**) for T cells from Wolf *et al.*^24^ for *n* = 2,610 proteins. (**c**) Activated versus resting protein clearance half lives in T cells. (**d**) Change in protein abundance, log_2_(activated/resting), versus change in protein clearance rate, log_2_(activated/resting), in T cells (Wolf *et al.*); *n* = 2,610.

**Fig. EV4.**
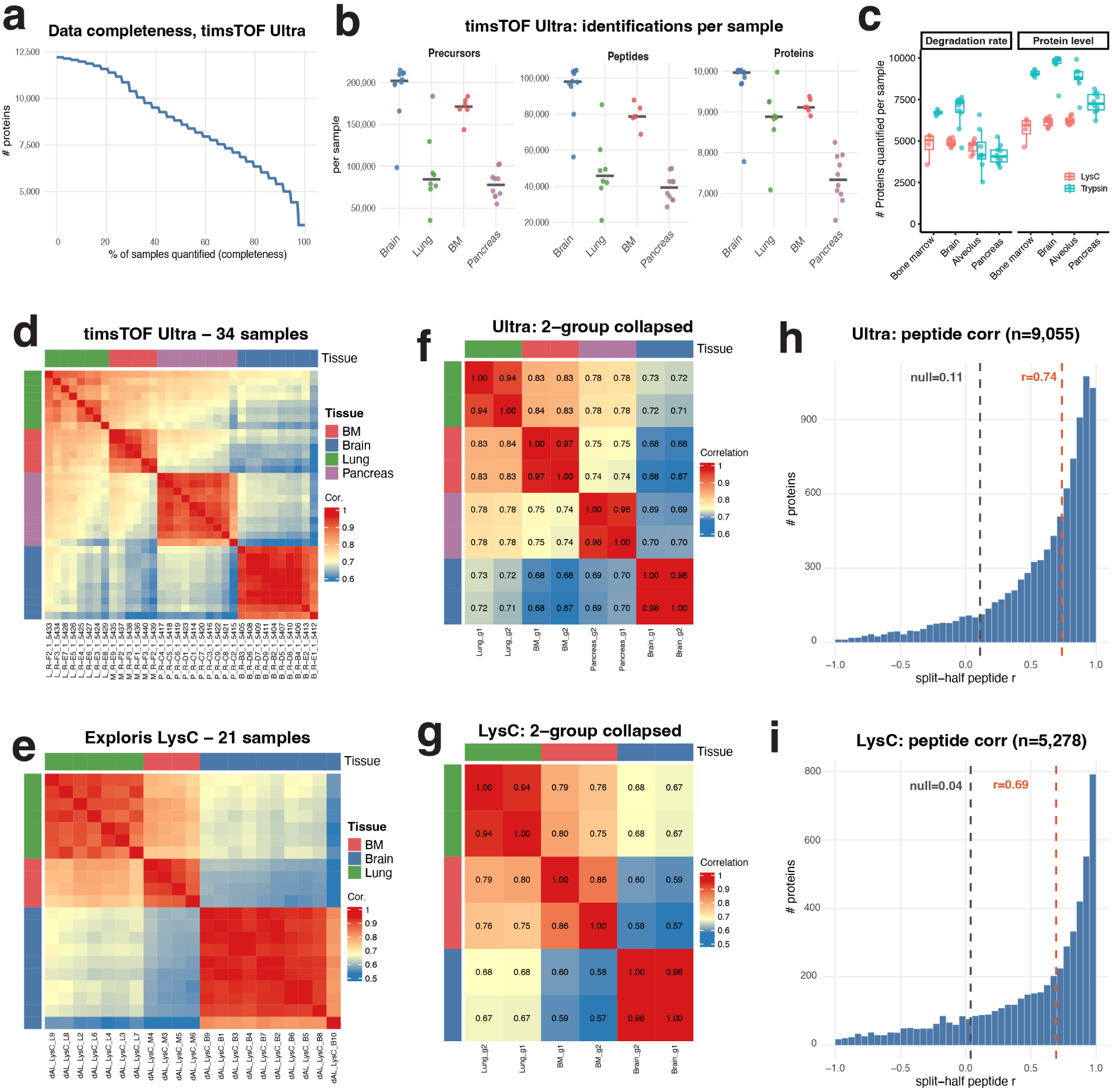
Within data set reproducibility and data quality. (**a**) Data completeness for the timsTOF Ultra dataset: number of protein groups (y-axis) quantified in at least a given fraction of samples (x-axis). (**b**) Number of identifications per timsTOF Ultra sample, separated by tissue, for precursors (heavy and light channels counted separately), unique peptide sequences, and protein groups. Each point is one sample; horizontal bars denote the per-tissue median. (**c**) Number of proteins with quantified degradation rates and protein-level abundances per sample, separated by tissue, for the Lys-C digest (Exploris 480) and trypsin digest (timsTOF Ultra). (**d**),(**e**) Pairwise Pearson correlations of protein abundances between all samples, for the (**d**) timsTOF Ultra and (**e**) Exploris 480 Lys-C datasets; top and side color bars indicate tissue. (**f**),(**g**) Pairwise correlations between eight pseudo-samples formed by splitting each tissue into two halves and averaging (2-group collapsed), for the (**f**) timsTOF Ultra and (**g**) Lys-C datasets. (**h**),(**i**) Distribution of split-half peptide correlations for the (**h**) timsTOF Ultra (*n* = 9,055 protein groups) and (**i**) Lys-C (*n* = 5,278) datasets; the grey dashed line marks the median of a shuffled peptide-to-protein null distribution.

**Fig. EV5.**
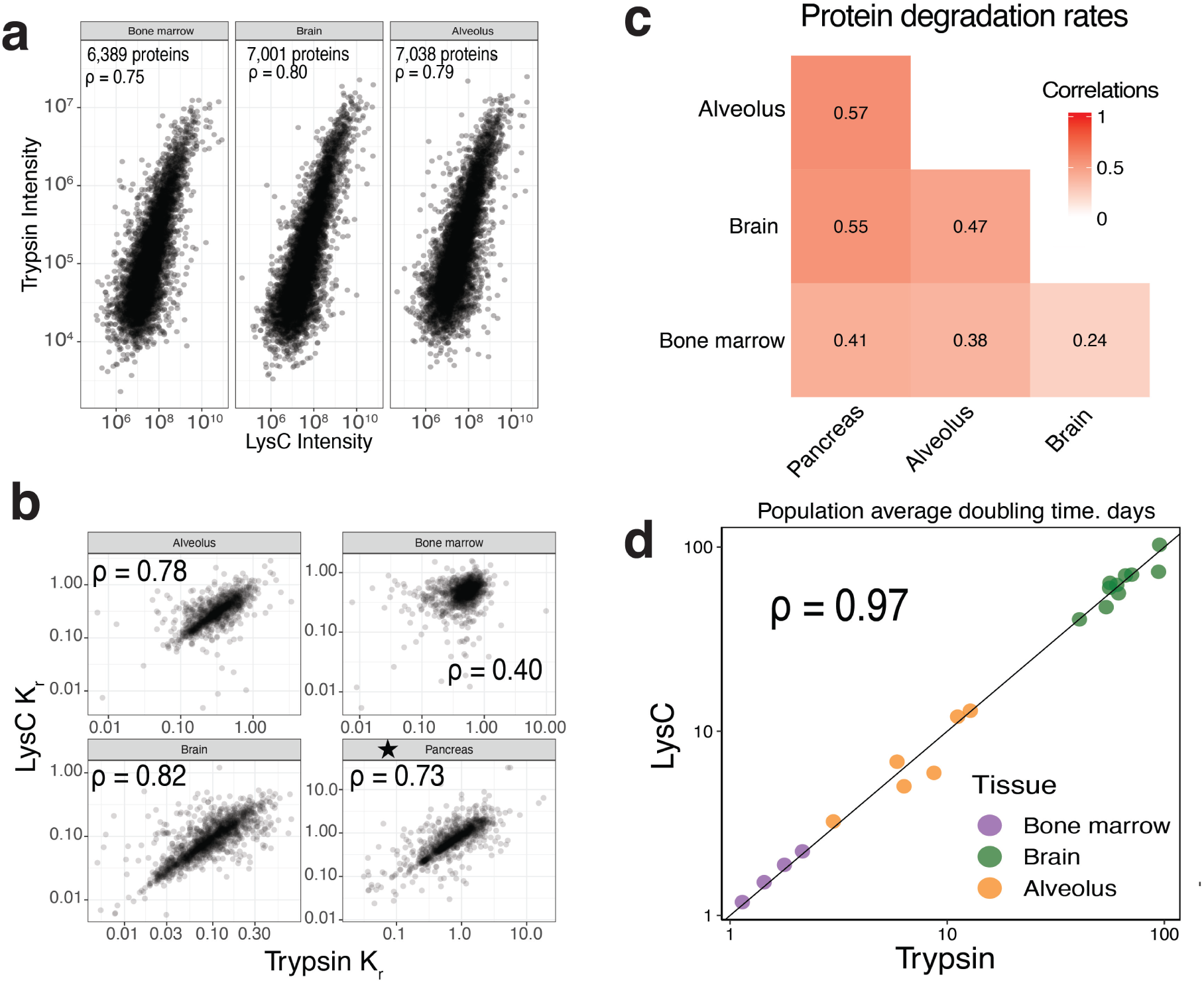
Quantitative accuracy and reproducibility of protein abundance and clearance rates for mouse tissues. (**a**) Reproducibility for protein abundance measurements for trypsin and Lys-C digests. (**b**) Reproducibility for clearance rate measurements for trypsin and Lys-C digests. Because the pancreas did not have a Lys-C condition due to the large amount of trypsin expressed in the tissue, technical replicates are plotted. (**c**) Correlation between average clearance rate measurements across tissues. (**d**) Reproducibility for measurements of cell doubling time as defined by average clearance rate of histones H3 and H4.

**Fig. EV6.**
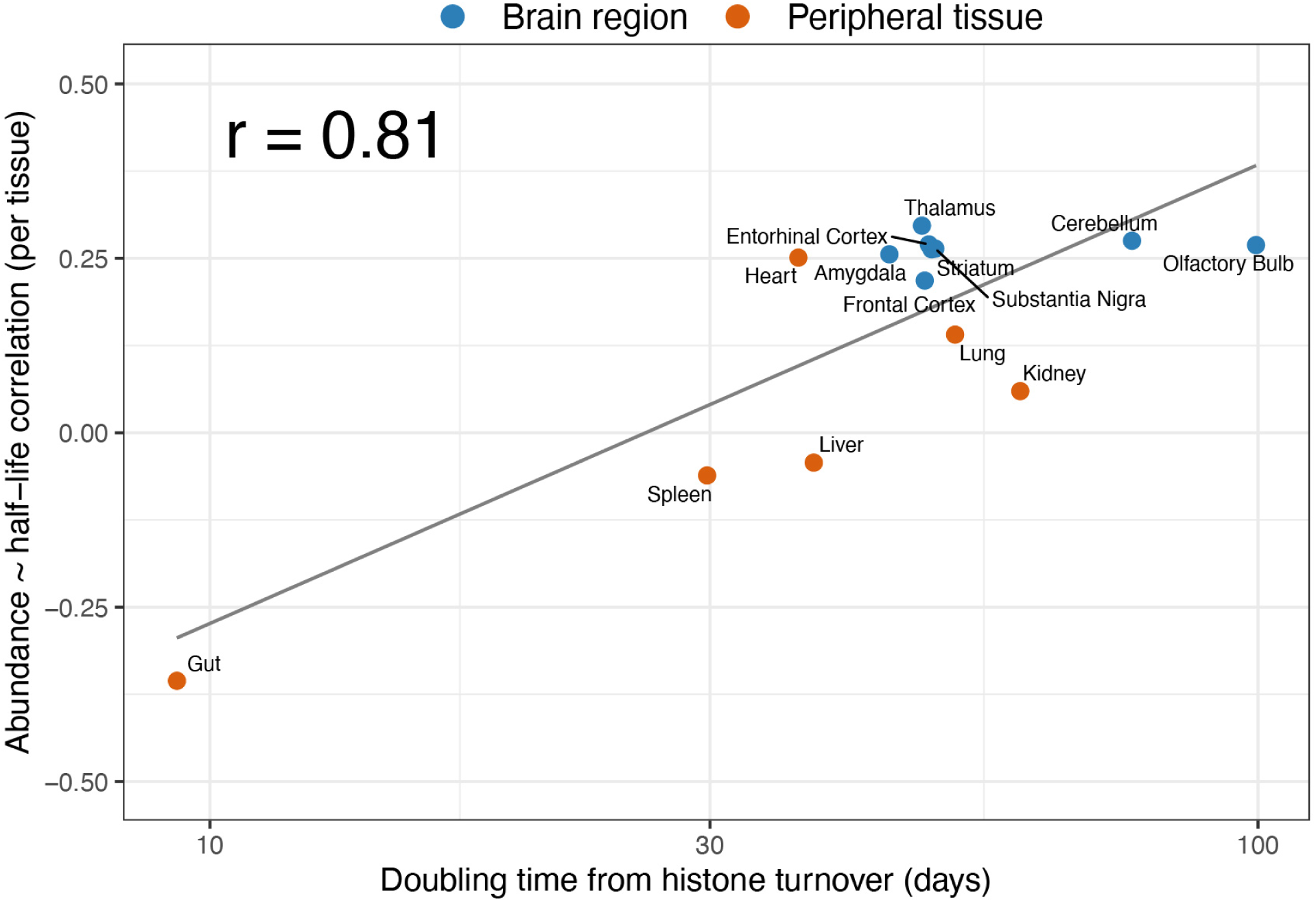
Contribution of protein half-life to abundance across mouse tissues. Each point is one tissue from the turnover atlas of Li *et al.* (8 brain regions, blue; 6 peripheral tissues, orange; 14 tissues total). The y-axis is the per-tissue correlation, across proteins, between protein abundance (iBAQ) and protein half-life; the x-axis is the tissue doubling time estimated from histone turnover (median histone half-life). Correlations and half-lives are computed on the set of proteins quantified in all tissues.

